# Standardising and harmonising research data policy in scholarly publishing

**DOI:** 10.1101/122929

**Authors:** Iain Hrynaszkiewicz, Aliaksandr Birukou, Mathias Astell, Sowmya Swaminathan, Amye Kenall, Varsha Khodiyar

**Affiliations:** Head of Data Publishing, Springer Nature; Senior Editor, Computer Science, Springer-Verlag GmbH; Marketing Manager, Nature Research; Head of Editorial Policy, Nature Research; Global Head of Life Sciences, Springer Nature; Data Curation Editor, *Scientific Data*

## Practice paper

Research data policies influence researchers’ willingness to share research data to varying extents (Meadows, 2014; Schmidt, Gemeinholzer, & Treloar, 2016). A growing number of research funders and institutions are introducing policies on research data sharing. These include the National Institutes of Health (NIH), Gates Foundation, the EU Horizon 2020 programme, Wellcome Trust and the seven UK research councils (Hahnel, 2015). Policy requirements vary, with some requiring researchers to prepare data management plans and others, such as the Engineering and Physical Sciences Research Council (EPSRC), requiring evidence of public data archiving to be included in published research papers. To support publication of more reproducible research scholarly journals, societies and conferences^1^ are also introducing data sharing policies which, in principle, should reflect the needs and norms of their respective research communities while being cognizant of funder requirements, where applicable.

But with thousands of journals across many different publishers, the research data policy landscape of journals is too complex (Naughton & Kernohan, 2016). Moreover, many journals have no stated policy on research data, and journals and publishers arguably have a responsibility to support researcher compliance with funder policies - as is established for open access policies (for papers, rather than data). As well as compliance with policy, enabling data sharing linked to scholarly publications can benefit individual researchers through increased visibility and citations (Piwowar & Vision, 2013), enable publishers to disseminate richer content, and also improve access to (and understanding of) data for further research.

An attempt by JISC to create a database of all journal research data policies - to complement previous projects cataloguing open access policies - was not completed in part due to the lack of standardisation and harmonisation of data policies across journals and communities (Naughton & Kernohan, 2016). The mandatory and optional aspects of stated policies on journal websites can often be ambiguous, and how these policies are enforced is also often unclear. Data sharing policies, ultimately, intend to promote the practice and publication of more open research. Open research data is an enabler of high quality research and innovation, as demonstrated in communities such as crystallography, genetics, archaeology and linguistics (*Concordat on Open Research Data*, 2016).

To address the complexities researchers face during publication, and the potential community-wide benefits of wider adoption of clear data policies, the publisher Springer Nature has developed a standardised, common framework for the research data policies of its more than 2,500 journals. An expert working group was convened (including IH, AB, SS AK and VK) to audit and identify common features of research data policies of the journals published by Springer Nature, where policies were present. The group then consulted with approximately 30 editors, covering all research disciplines, within the organisation. The group also consulted with academic editors and librarians and funders, which informed development of the framework and the creation of supporting resources, such as Frequently Asked Questions (http://www.springernature.com/gp/group/data-policy/faq) Four types of data policy were defined (Figure 1).

**[Figure 1]:**
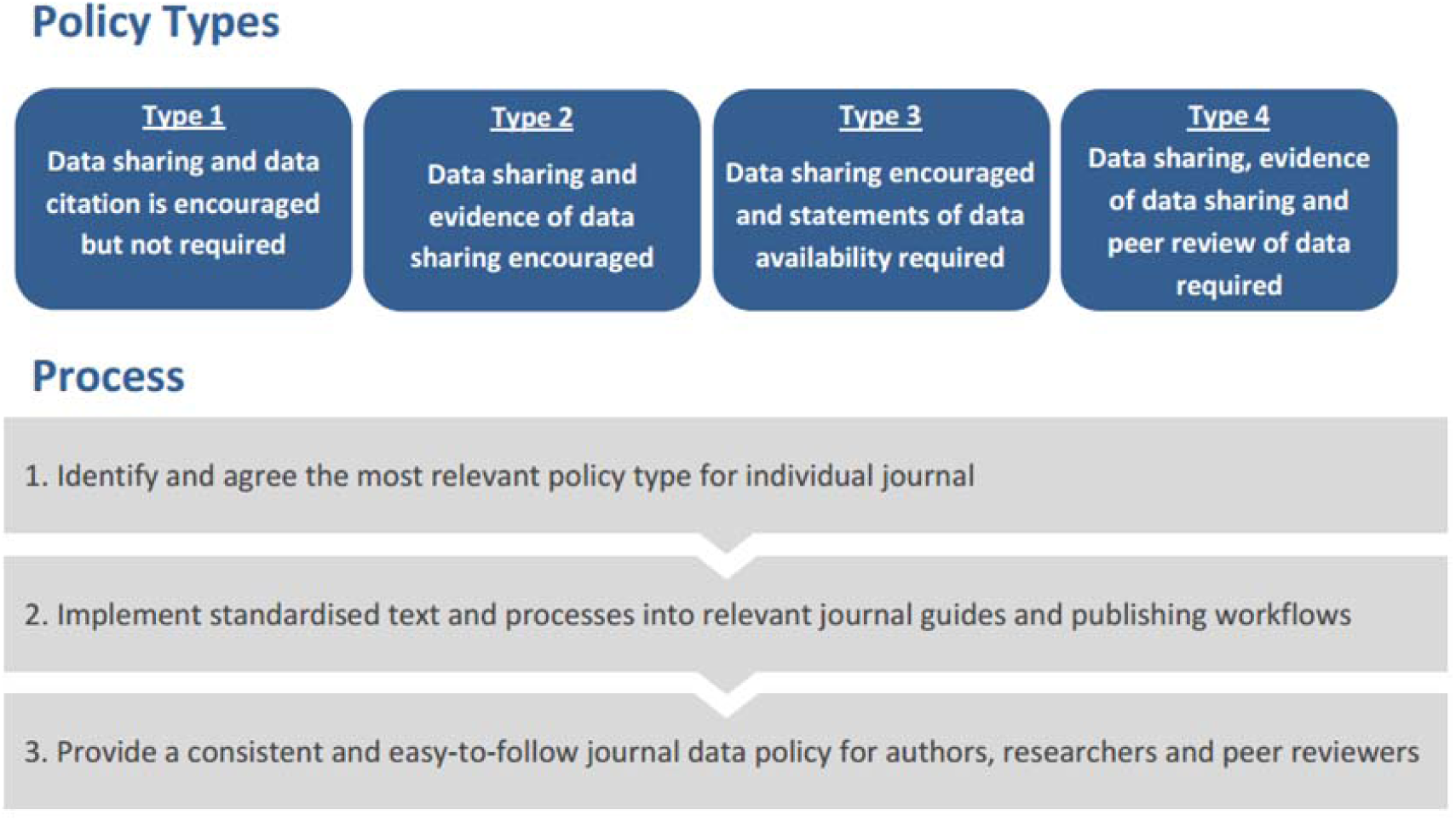
Overview of the four data policy types.

Individual journals and research communities are at different stages of readiness to support data sharing, from communities that may mandate Open Data to those who are just beginning to debate the issues. Defining four types of policy is intended to recognise these differences whilst providing a common and easy-to-understand framework to encourage good, and better, data sharing practices across all research areas.

The Springer Nature research data policy framework divides data policy for publications into nine features (Figure 2), and each type of policy has a defined number of features, with the type one policy having the fewest features and the type four policy the most (Figure 2). The type one policy encourages data sharing and data citation and provides researchers with a list of data repositories, and the type two policy provides information on preparing data availability statements. There is increased expectation for compliance with the type 3 policy, which requires data availability statements. The type four policy requires open data and requires peer reviewers to access data supporting publications. The policy texts are available in full under a Creative Commons licence at: http://www.springernature.com/gp/group/data-policy/policy-types

**[Table 1].** The policy types and their features.

Key: ● = Mandatory ◐ = Optional ○ = Not Required
| Feature | Explanation | Type 1 | Type 2 | Type 3 | Type 4 |
| --- | --- | --- | --- | --- | --- |
| **Data sharing via repositories supported** | Details of sharing via repositories is referred to in journal guide to authors | ● | ● | ● | ● |
| **Data citation permitted** | Journal style guide permits authors to cite publicly available datasets in reference lists | ● | ● | ● | ● |
| **Publisher helpdesk** | Helpdesk contact details included in journal information for authors | ◐ | ◐ | ◐ | ◐ |
| **Public data deposition and dataset identifier checks for specific types of data** | Data deposition checked as part of the publishing process where there is an established research community mandate | ○ | ◐ | ● | ● |
| **Data availability statements** | Statement in published articles explaining how supporting data can be accessed | ○ | ◐ | ● | ● |
| **Public data deposition and dataset identifier required and verified** | Data made publicly available and data identifiers provided for all published articles (with exceptions for sensitive/personal data) | ○ | ○ | ◐ | ● |
| **Data citations** | Relevant dataset citations in reference lists provided and verified | ○ | ○ | ◐ | ● |
| **Peer review of data** | Peer reviewer guidelines and process give guidance on accessing and reviewing data files | ○ | ○ | ◐ | ● |
| **Integrated data repository** | Submission system/review process integrated with a journal-specific or general repository, such as figshare | ○ | ○ | ◐ | ● |

Importantly, the type of data sharing policy a journal introduces is not determined by journal quality, prestige or Impact Factor. Journals adopt the data sharing policy that is most appropriate to its research community and the resources available to that community-encouraging the most relevant good practice for their community.

**[Table 2].** Examples of journals following each type of data policy.

| Policy type | Example journal | Web link |
| --- | --- | --- |
| 1 | *Machine Vision and Applications* | See ‘Instructions for Authors’ |
| 2 | *Plant and Soil* | See ‘Instructions for Authors’ |
| 3 | *Palgrave Communications* | See 'Editorial policies' |
| 4 | *Scientific Data* | See 'Data policies' |

Our initial review of existing data policies in Springer Nature journals found that life science journals – Nature and BioMed Central journals in particular – were more likely to have some form of data policy. However, the policy framework was developed agnostic of research disciplines and has been adopted by journals covering many disciplines including the humanities, social sciences engineering, physical and computer sciences and mathematics.

Repository information, data citations and helpdesk support are features of all four types of policy. The helpdesk (http://www.springernature.com/gp/group/data-policy/helpdesk) is intended to advise researchers on complying with data policies and on finding suitable repositories. It is also intended to support Editors in identifying and implementing an appropriate research data policy.

The policies have begun to be introduced to Springer Nature journals, which include the Nature titles, BioMed Central journals, and those hosted on SpringerLink, in the second quarter of 2016 (Hrynaszkiewicz, 2016). As of January 2017 more than 700 journals have adopted a standard policy and this number is growing weekly. As of January 2017, the type 3 policy is the most common policy type having been adopted by more than 340 journals. This may be biased by BioMed Central journals such as *BMC Genomics* and Nature journals all largely adopting this type of policy (“Announcement: Where are the data?,” 2016). A list of all journals and the type of policy they have adopted is available at http://www.springernature.com/gp/group/data-policy/journal-listing. While there is a disciplinary trend towards adoption of the type 2 or 3 policy by life science journals, other factors also influence policy type selection. Journals with fewer editorial office staff to perform compliance checks, for example, may be more likely to select the type 1 or 2 policy, and journals with no previous data sharing policy are also more likely to adopt the type 1 or 2 policy. The type 4 policy, requiring open data for every publication, is the least common and has so far only been adopted by data journals with either a strong focus on data sharing and/or an author community with well-established cultures and repositories for open data sharing. There are plans to introduce data policies to books and conference proceedings that report original research.

In its first six months of operation the helpdesk was contacted by authors as well as professional and academic editors. The most common questions related to finding data repositories, choosing and implementing a policy type, and preparing data availability statements. Authors can also be advised on funders’ policies and other aspects of data sharing through the helpdesk.

Resources for authors and editors are provided via the publisher’s website, including a list of repositories (http://www.springernature.com/gp/group/data-policy/repositories) and data availability statement guidance (http://www.springernature.com/gp/group/data-policy/data-availability-statements), but individual journals’ guides to authors are also updated when a policy is introduced. Journal and community policy requirements have been found to be stronger incentives for researchers to share data than publisher policy (Schmidt et al., 2016).

Further studies exploring the costs and benefits of data policy implementation at the different levels are currently being conducted and planned. For example, measuring the additional time taken by editors to require data availability statements in published papers (preliminary data reported in: http://www.stm-assoc.org/2016_12_06_Digital_Publishing_Hrynaszkiewicz_Research_data.pdf). Providing editors and other stakeholders with evidence of this kind will further support informed decision making about policy adoption and implementation.

To potentially enable standardisation and harmonisation of data policy across funders, institutions, repositories, societies and other publishers the policy framework was made available under a Creative Commons license (“Over 600 Springer Nature journals commit to new data sharing policies,” n.d.) for reuse by other organisations. However, the framework requires wider debate with these stakeholders and we plan to work with the Research Data Alliance (RDA) to initiate this process (https://www.rd-alliance.org/groups/data-policy-standardisation-and-implementation).

## Footnotes

1 see ECML PKDD 2016 call for papers at http://www.ecmlpkdd2016.org/submission.html#call-conference or ISWC 2016 call for resource papers at http://iswc2016.semanticweb.org/pages/calls/resource-track.html.

## Acknowledgements

For contributions to policy and repository list development, Dr Andrew Hufton, Managing Editor, *Scientific Data*.

This paper was presented at the 12th International Digital Curation Conference, Edinburgh, UK on 22 February 2017 (http://www.dcc.ac.uk/events/idcc17/programme) and will be submitted to International Journal of Digital Curation.

